# ACValidator: a novel assembly-based approach for *in silico* validation of circular RNAs

**DOI:** 10.1101/556597

**Authors:** Shobana Sekar, Philipp Geiger, Jonathan Adkins, Geidy Serrano, Thomas G. Beach, Winnie S. Liang

**Affiliations:** Neurogenomics division, Translational Genomics Research Institute, Phoenix, Arizona, United States of America; Arizona Alzheimer’s Consortium, Phoenix, Arizona, United States of America; Banner Sun Health Research Institute, Sun City, Arizona, United States of America

## Abstract

Circular RNAs (circRNAs) are evolutionarily conserved RNA species that are formed when exons ‘back-splice’ to each other. Current computational algorithms to detect these back-splicing junctions produce divergent results, and hence there is a need for a method to distinguish true positive circRNAs. To this end, we developed ACValidator (<u>A</u>ssembly based <u>C</u>ircRNA <u>Validator</u>) for *in silico* validation of circRNAs. ACValidator extracts reads from a user-defined window on either side of the circRNA junction and assembles them to generate contigs. These contigs are aligned against the circRNA sequence to find contigs spanning the backspliced junction. When evaluated on simulated datasets, ACValidator achieved over 80% sensitivity and specificity on datasets with an average of 10 circRNA-supporting reads and with read lengths of at least 100 bp. In experimental datasets, ACValidator produced higher validation percentages for samples treated with ribonuclease R compared to non-treated samples. Our workflow is applicable to non-polyA-selected RNAseq datasets and can also be used as a candidate selection strategy for experimental validations. All workflow scripts are freely accessible on our github page https://github.com/tgen/ACValidator along with detailed instructions to set up and run ACValidator.

**Author summary:** Circular RNAs (circRNAs) are a recent addition to the class of non-coding RNAs and are produced when exons ‘back-splice’ and form closed circular loops. Although several computational algorithms have been developed to detect circRNAs from RNA sequencing (RNAseq) data, they produce divergent results. We hence developed the software <u>A</u>ssembly based <u>C</u>ircular RNA <u>Validator</u> (ACValidator) as an orthogonal strategy to separately validate predicted circRNAs *in silico*. ACValidator takes as input a sequence alignment mapping (SAM) file and the circRNA coordinate(s) to be validated. Reads surrounding the circRNA junction are extracted from the SAM file and assembled to generate contigs. These contigs are then aligned against the circRNA sequence to identify contigs that span the back-spliced junction. We evaluated our workflow on simulated as well as experimental datasets to demonstrate the utility of our approach. ACValidator is implemented in python and is highly computationally efficient, with a run time of less than 2 minutes for an 8 GB SAM file. This workflow is applicable to non-polyA-selected RNAseq datasets and can also be used as a candidate selection strategy for experimental validations.

## Introduction

Circular RNAs (circRNAs) represent a large class of ubiquitously expressed non-coding RNAs that are formed when exons ‘back-splice’ to each other. The advent of high throughput RNA sequencing (RNAseq) technologies and bioinformatics algorithms has facilitated the identification of thousands of circRNAs in multiple cell and tissue types [1-4]. These studies have found that circRNAs are highly abundant and evolutionarily conserved, as well as exhibit cell typeand developmental stage-specific expression. CircRNAs are also more stable than linear RNAs since they are covalently closed loops without 5’/3’ termini or a polyadenylated tail. Furthermore, studies investigating their functional relevance have revealed that circRNAs can act as miRNA regulators [3,5-7], decoys to RNA binding proteins [8], and regulators of parental gene transcription [9].

Several computational tools have been developed to identify these back-splicing events in RNAseq data. Strategies employed by these computational tools to identify circRNAs include: 1) a pseudo-reference-based strategy, which is used by KNIFE [10]; and 2) a fragment-based strategy, which is used by find_circ [3], CIRCexplorer [11], Mapsplice [12] and DCC [13]. While KNIFE constructs a pseudo-reference of all possible out-of-order exons to align reads against, fragment-based strategies detect circRNAs based on the mapping information of a split read’s alignment to the reference genome [14]. When segments of a split read align to the reference in a non-colinear order, they are marked as potential circRNA candidates. Apart from these strategies, CIRI uses CIGAR (concise idiosyncratic gapped alignment report) signatures in the alignment file to identify circRNAs [15].

Tool comparison studies have revealed that existing circRNA detection algorithms produce divergent results due to the use of different aligners, heuristics and filtering criteria [16,17]. Hence, there is a need for an *in silico* validation approach that can distinguish true versus false positive circRNAs identified using these algorithms. To this end, we developed ACValidator (<u>A</u>ssembly based <u>C</u>ircular RNA <u>Validator</u>), which can be used as an *in silico* validation strategy as well as a candidate selection tool for experimental validation. While existing approaches focus on detection of circRNAs, our approach performs *in silico* validation of circRNAs detected using these existing approaches. ACValidator first extracts reads from a fixed window on either side of the circRNA junction of interest from the alignment file and assembles them to generate contigs. These contigs are then evaluated for alignment against the circRNA junction sequence. We defined four different stringency criteria, ranging from 10 to 60 base pairs (bp) overlap across the junction, to capture as many validations as possible. When evaluated on simulated as well as experimental datasets, ACValidator achieves higher precision and sensitivity in datasets with higher circRNA coverage compared to ones with lower coverage.

## Design and Implementation

ACValidator takes as input a sequence alignment mapping (SAM) file and the circRNA coordinate(s) to be validated (Fig 1). ACValidator operates in three phases: (1) extraction and assembly of reads from the SAM file to generate contigs (2) generation of a pseudo-reference file, and (3) alignment of contigs from phase 1 against the pseudo-reference from phase 2. First, reads are extracted from a user-defined window *w* on either side of the given SAM file [(*start-coordinate + w)* and (*end-coordinate w*); where start-coordinate is the splice acceptor and end-coordinate is the splice donor of the circRNA junction]. Our datasets were run using two different window sizes, where *w* = insert size or *w* = 2 * insert size, in order to capture as many reads overlapping the circRNA junction as possible, and to understand the effect of window size on the results. Users can adjust this parameter based on their library insert size or read length. The tool thus extracts aligned reads within *w* bp on either side of the junction from the SAM file using SAMtools [18]. The extracted reads are then converted into FASTQs and assembled using the Trinity assembler [19] to generate contigs (FASTA file). In the second phase, a pseudo-reference of the sequence surrounding the circRNA junction of interest is generated. This is performed by also extracting *w* bp from the end and start of the circRNA junction from the genome reference FASTA file (GRCh37) and concatenating the two sequences from end to end to capture the sequence on either side of the circular junction. Lastly, the assembled contigs from phase 1 are aligned to the pseudo-reference from phase 2 using the widely adapted BWA-MEM [20] aligner. Each resulting alignment record is then examined to check whether it overlaps with the circRNA junction sequence using four different stringency criteria. The criteria require the following minimum lengths of alignment on both sides of the circRNA junction: high-stringency—30 bp (total 60 bp overlap); medium-stringency—20 bp (total 40 bp overlap); low-stringency—10 bp (total 20 bp overlap); and very-low-stringency—5 bp (total 10 bp overlap). These stringency cut-offs were defined in order to capture as many validations as possible while still accounting for the extent of overlap between the contig and the circRNA junction sequence, as well as to assess whether we observe differences in the sensitivity measurements across these cut-offs.

**Fig 1.**
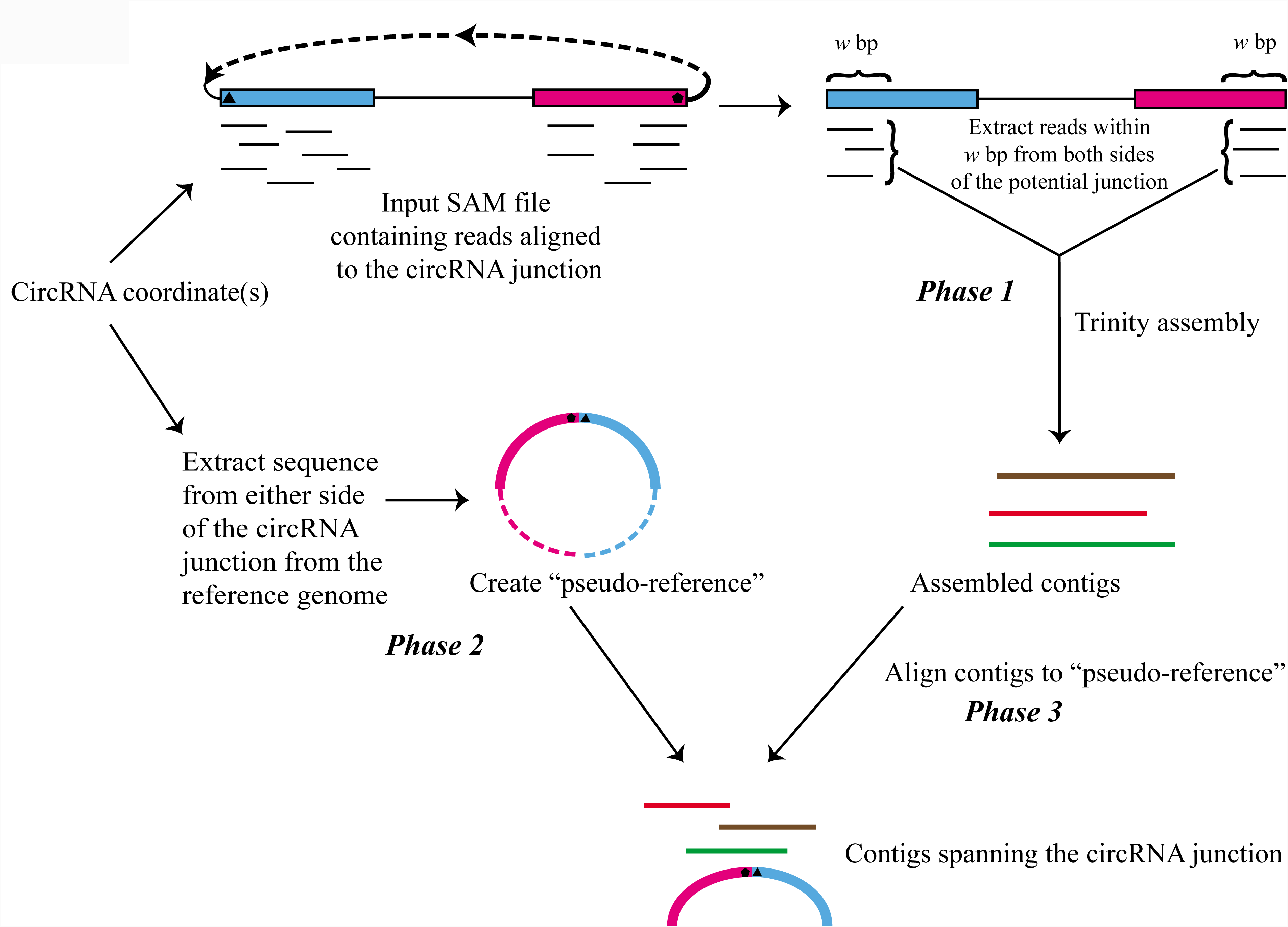
ACValidator workflow. ACValidator takes as input the sequence alignment mapping (SAM) file and the circRNA junction coordinate(s) to be validated. In phase 1, reads from either side of the junction within a user-defined window (*w*) are extracted and assembled using Trinity. A pseudo-reference containing the circRNA sequence of interest is generated from the reference genome in phase 2. The pseudo-reference consists of *w* bp from either side of the circRNA junction of interest (solid blue and pink blocks in phase 2). The broken blue and pink segments represent the remaining portions of the exons that constitute the circRNA but they are not part of the pseudo-reference. Lastly, in phase 3, the assembled contigs are aligned to this pseudo-reference and checked for overlap with the sequence of the junction to be validated.

### Datasets used for evaluation

#### Simulated datasets

We used CIRI-simulator [15] to generate nine synthetic RNAseq datasets that had variable average number of supporting reads for circular and linear RNAs (2-10), as well as three different read lengths (50, 100, 150bp) to evaluate the performance of our workflow (Table 1). CIRI-simulator takes a FASTA-formatted reference file and a GTF annotation file as input, and generates circular and linear RNA sequences. Recently, Zeng *et al*. [17] re-designed this tool to generate synthetic reads for circRNAs deposited in circBase [21]. We generated simulated datasets with minimum circRNA size of at least 50 bp and insert size of 300 bp. Overall, an average of 89,293 circRNAs were generated across these nine simulated datasets. CIRI-simulator ensures these circRNAs map to locations distributed across the entire genome, thereby eliminating any bias associated with genomic location (Fig 2). The generated true positive simulation datasets are named using the convention pos_<circRNA_coverage>_<linearRNA_coverage>_ <read_length> (pos: positive; Table 1).

**Table 1.** Simulation dataset parameters. All simulation datasets based on data generated from human cerebellum and diencephalon, SH-SY5Y cells, Hs68 cells, HeLa cells and HEK293 cells. Columns 3, 5, 6 and 7 are user-defined parameters supplied to CIRI-simulator. Minimum circRNA size used in simulation: 50 bp.

| Simulation set | Simulation dataset name | Average # of circRNA support-ing reads | Range of # of circRNA support-ing reads | Average # of linear RNA sup- porting reads | Read length (bp) | Insert size (bp) | # of reads generated | Window lengths tested (bp) | Overlapping base thresholds tested (bp) |
| --- | --- | --- | --- | --- | --- | --- | --- | --- | --- |
| 1 | pos_10_2_150 | 10 | 3 to 27 | 2 | 150 | 300 | 18,768,200 | 300, 600 | 60, 40, 20, 10 |
| 2 | pos_10_2_100 | 10 | 2 to 25 | 2 | 100 | 300 | 28,024,890 | 300, 600 | 60, 40, 20, 10 |
| 3 | pos_10_2_50 | 10 | 2 to 26 | 2 | 50 | 300 | 55,794,356 | 300, 600 | 60, 40, 20, 10 |
| 4 | pos_5_5_150 | 5 | 2 to 17 | 5 | 150 | 300 | 10,867,166 | 300, 600 | 60, 40, 20, 10 |
| 5 | pos_5_5_100 | 5 | 2 to 16 | 5 | 100 | 300 | 16,173,002 | 300, 600 | 60, 40, 20, 10 |
| 6 | pos_5_5_50 | 5 | 2 to 17 | 5 | 50 | 300 | 32,090,962 | 300, 600 | 60, 40, 20, 10 |
| 7 | pos_2_10_150 | 2 | 2 to 10 | 10 | 150 | 300 | 7,209,692 | 300, 600 | 60, 40, 20, 10 |
| 8 | pos_2_10_100 | 2 | 2 to 12 | 10 | 100 | 300 | 10,688,368 | 300, 600 | 60, 40, 20, 10 |
| 9 | pos_2_10_50 | 2 | 2 to 11 | 10 | 50 | 300 | 21,121,578 | 300, 600 | 60, 40, 20, 10 |

Pos: positive

**Fig 2.**
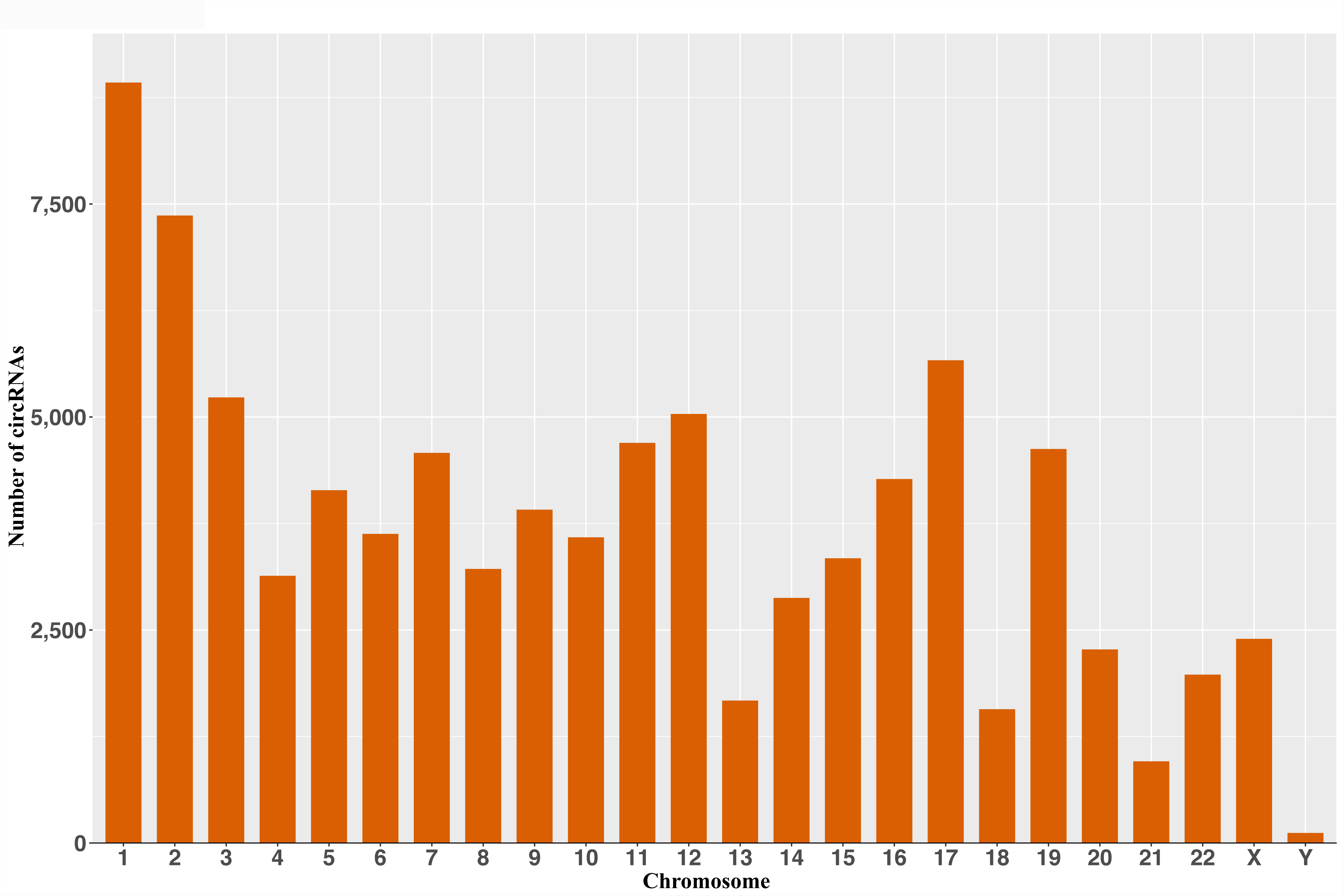
Chromosomal distribution of simulated datasets. Simulation datasets were generated using CIRIsimulator, which ensures circRNAs are simulated across all chromosomes. The distribution of generated circRNAs across the different chromosomes is similar to the chromosomal size distribution.

#### Experimental datasets

To test ACValidator on experimental data, we analyzed six pairs of ribonuclease R (RNase R)-treated and non-treated samples (N = 12, Table 2). RNase R is an exoribonuclease that selectively digests linear RNA but leaves behind circular structures, and it is hence widely used for circRNA enrichment. Three of these sample pairs were downloaded from the sequence read archive (SRA) [22] and were generated from HeLa and Hs68 cell lines treated or not treated with RNase R. The remaining three sample pairs were generated in-house from total RNA extracted from the middle temporal gyrus (MG) of three human healthy elderly control brains (manuscript under preparation). All data generated through this study are accessible through the European Genome Archive (EGA; accession EGAS00001003128).

**Table 2.**
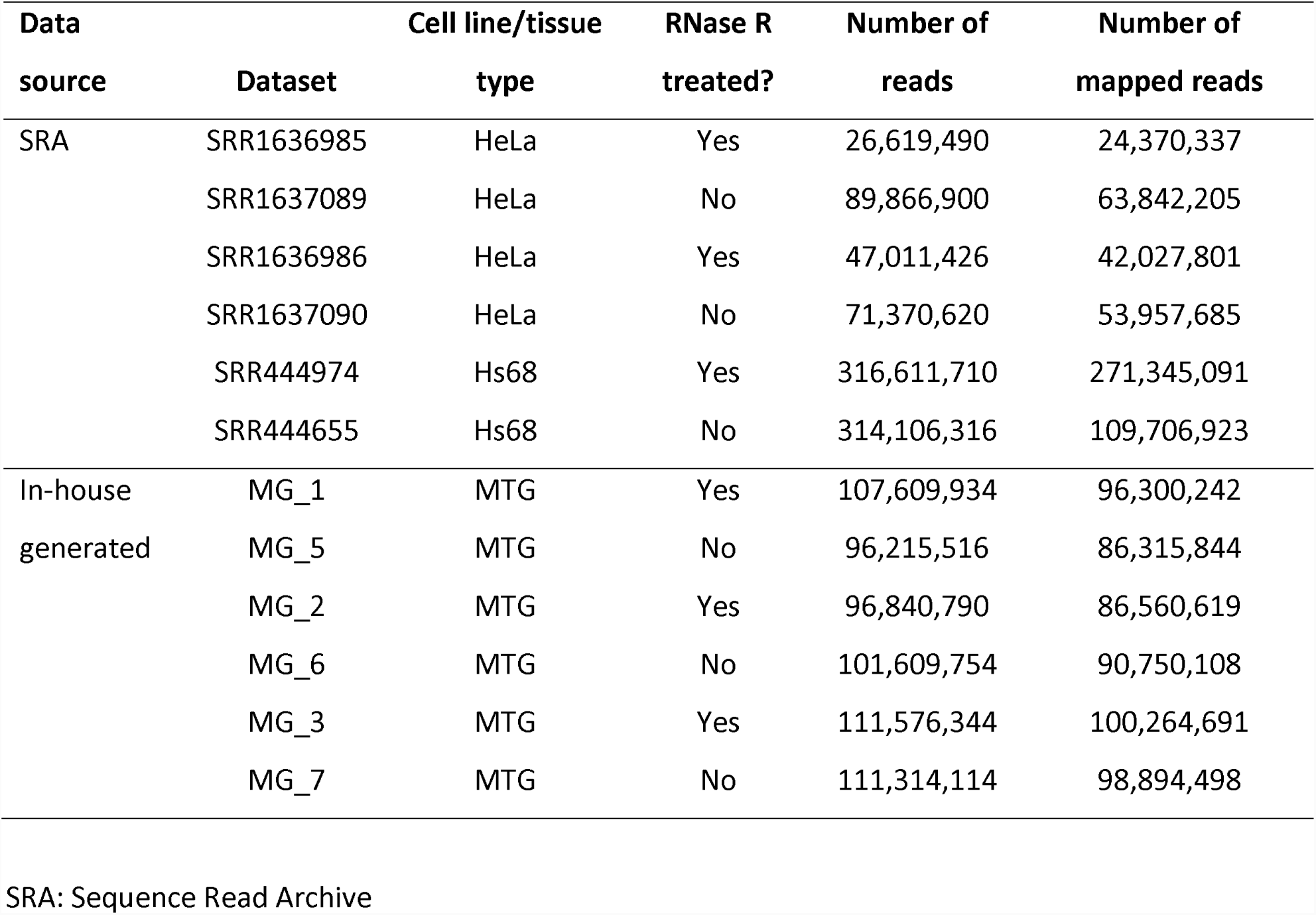
Summary of experimental, non-simulated datasets used in this study.

#### Polymerase chain reaction validation

Polymerase chain reaction (PCR) was performed to experimentally validate the presence of five most highly expressed circRNA candidates that were *in silico* validated by ACValidator. cDNA was synthesized from 500 ng of RNA isolated from the MG of the three tissue samples of interest using SuperScript II reverse transcriptase (ThermoFisher Scientific, Waltham, MA). Primers were designed to target the flanking exons of selected circRNA junction sites using Primer3 (http://bioinfo.ut.ee/primer3-0.4.0/). PCR conditions were optimized for each primer set and product sizes were assessed on a TapeStation 4200 instrument (Agilent technologies, Santa Clara, CA).

#### Software requirements/dependencies

ACValidator can be implemented on a Linux-based high-performance computing cluster and has minimal requirements and dependencies. These requirements include the following: (a) Trinity v2.3.1 or above; (b) Python v2.7.13 or higher with the pysam package installed; (c) Bowtie2 v2.3.0 [23] or above; (d) SAMtools v1.4 or above and (e) BWA v0.7.12 or above.

## Results and discussion

### Performance evaluation of ACValidator using simulated data

To evaluate ACValidator, we generated nine simulation datasets with varying circular, linear RNA coverages and read lengths (Table 1). Since highly expressed circRNAs are more often of interest, we evaluated the top 100 most highly expressed circRNAs from each dataset, similar to a previous study [16]. We also selected 100 random non-circRNA coordinates as false positive candidates from each dataset (S1 Table). We replicated our analysis using two different window sizes: 1) *w* = insert size (300 bp) 2) *w* = 2 * insert size (600 bp) (section: Design and implementation).

We observed that simulations with higher circRNA coverages and longer read lengths achieved higher sensitivity (Fig 3). Specifically, when using *w* = 300 and overlap cut-off of 10 bp between the contig and pseudoreference, simulations 1 and 2, which have the highest circRNA coverage (10) and read lengths (150 bp, 100 bp) achieved the highest sensitivity of 90% and 89%, respectively. However, for simulation 3 where the read length was only 50 bp, the sensitivity fell to 38%. As the circRNA coverage decreases, the sensitivity also gradually reduces to 81% and below, with a further reduction in sensitivity to 46% and 28% observed for datasets with shorter read lengths (simulations 6 and 9). In datasets with lower circRNA coverage and/or shorter read length, this reduction in sensitivity was because Trinity did not find sufficient reads to assemble across these regions and hence was not able to generate contigs. We detect a similar pattern when using *w* = 600 and did not observe a drastic difference in sensitivity between the two window sizes (S1 Table). Further, we calculated the F1 score [F1 = (2 * Precision * Sensitivity)/ (Precision + Sensitivity)], a measure of accuracy, which indicates how well a tool achieves sensitivity and precision simultaneously (Fig 3). We found the results to be consistent with sensitivity measurements, indicating that higher circRNA coverage coupled with longer read length yields better performance of our approach.

**Fig 3.**
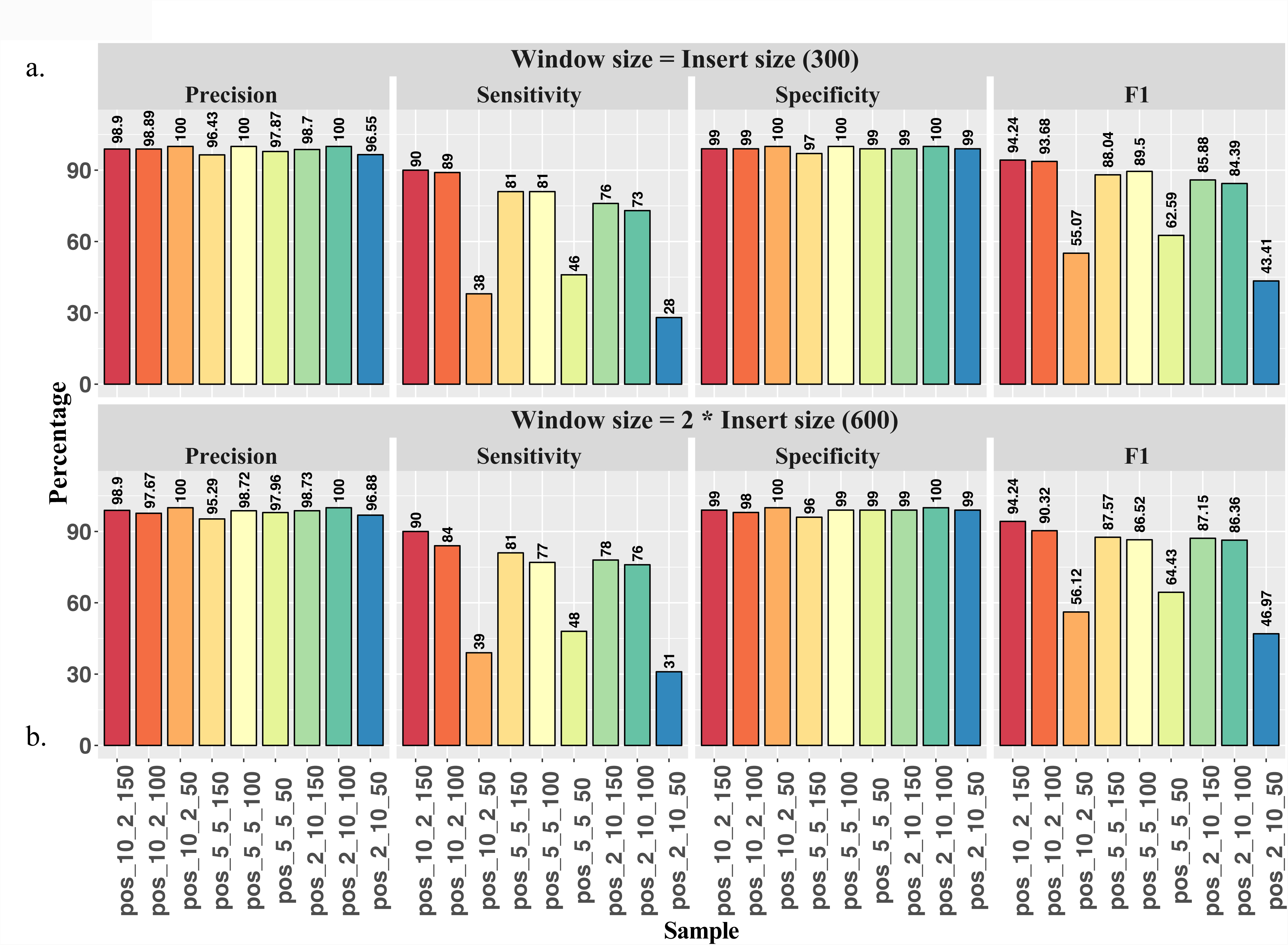
ACValidator performance on top 100 circRNAs and 100 non-circRNA candidates from simulated datasets. The 100 candidate circRNA junctions with the highest number of supporting reads were considered true positive (TP) and 100 randomly selected non-circRNA junction coordinates were considered true negative (TN). The simulation datasets are described in Table 1 and each simulation dataset (x-axis) is named using the naming convention: pos_<circRNA_coverage>_<linearRNA_coverage>_<read_length> (pos: positive). The panels represent ACValidator performance on top 100 TP candidates when using an overlap cut-off of 10 bp between the contig and pseudo-reference, and a) window size = insert size (300 bp) and b) window size = 2 * insert size (600 bp). P =TP/(TP+FP); S = TP/(TP + FN); Sp = TN/(TN+FP); F1 = (2 * P * S)/ (P + S). FP, false positives; FN, false negatives; S, sensitivity; Sp, specificity; P, precision.

We extended our analysis to the top 200 most highly expressed as well as bottom 200 least expressed candidates (S2, S3 Tables respectively), and observed a similar trend in performance (Fig 4). Additionally, we evaluated sensitivity using four stringency thresholds for the number of overlapping bases (30 bp, 20 bp, 10 bp and 5 bp; section: Design and implementation). For the top 200 highly expressed circRNAs, we achieved sensitivity of over 80% for simulation datasets 1 and 2, which had the highest circRNA coverage (10) and read lengths (150 and 100), using both window sizes (300 and 600). However, we did not observe a notable difference in the number of validations across the different thresholds for the number of overlapping bases, with an average of 86.8% validation for simulation set 1 (pos_10_2_150), 71.5% validation for simulation set 7 (pos_2_10_150) and 27.3% validation for simulation set 9 (pos_2_10_50; Fig 4a, S2 Table). For the bottom 200 least expressed candidates, lower stringency criteria resulted in more validations than the higher stringency criteria, with an average of 19.3% and 10.1% validation for the very low and high stringencies respectively. The overall validation percentages were below 45% across all the simulation datasets for these bottom 200 circRNA candidates (Fig 4b, S3 Table).

**Fig 4.**
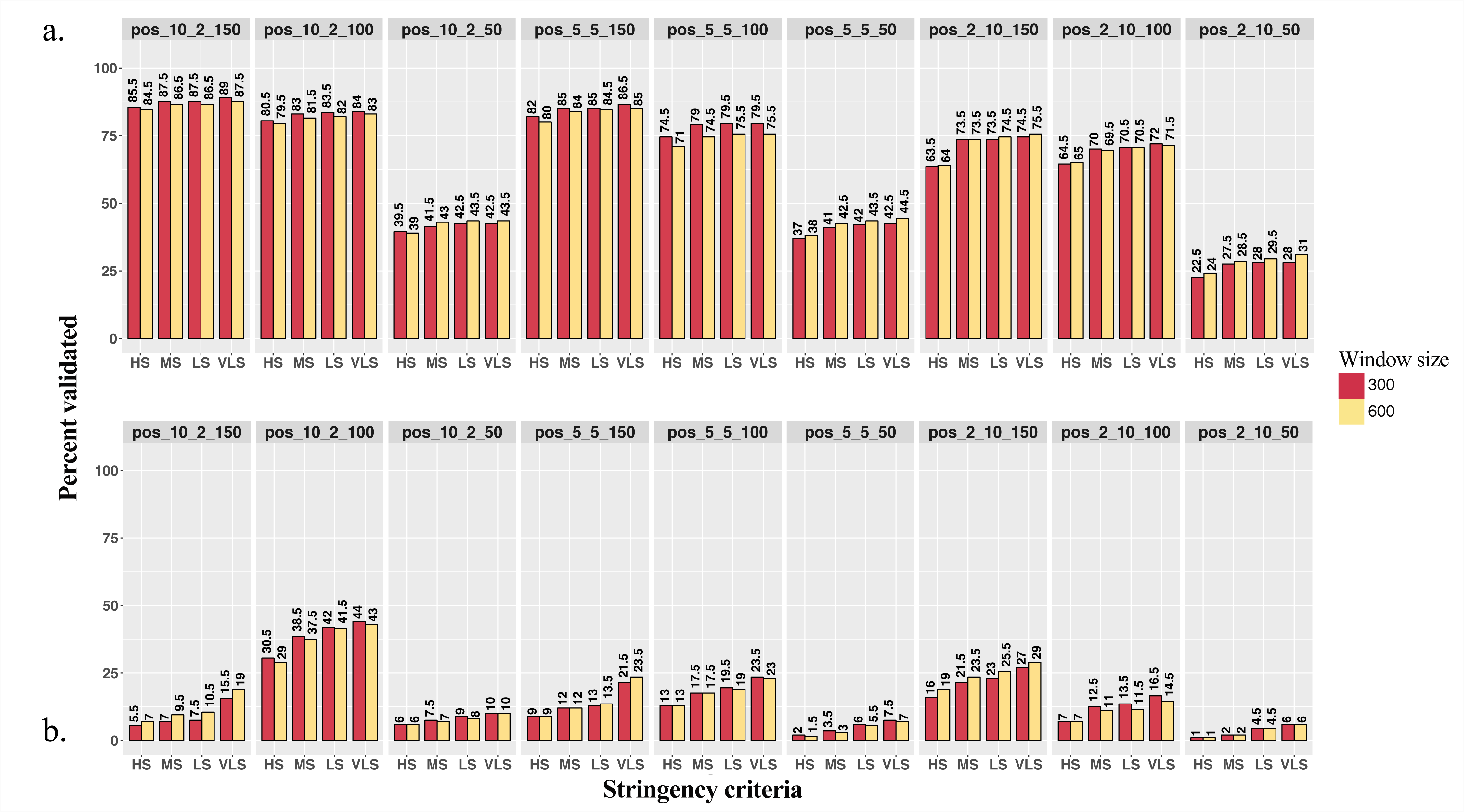
ACValidator performance on the top and bottom 200 circRNAs from simulated datasets. The (a) top and (b) bottom 200 candidates based on the number of circRNA supporting reads were evaluated using ACValidator, with two different window sizes, 300 and 600 bp. HS, high stringency; MS, medium stringency; LS, low stringency; VLS, very low stringency.

### Performance evaluation of ACValidator using experimental data

We next evaluated ACValidator using experimental, non-simulated datasets generated from human tissues or cell lines (Table 2). Since we do not know the true positive circRNAs for these datasets, we ran ACValidator on circRNAs that were called in both the RNase R-treated and non-treated datasets using six existing circRNA detection algorithms, find_circ, CIRI, Mapsplice, KNIFE, DCC and CIRCexplorer. Each tool was run using RNAseq aligners and parameter settings as recommended by the respective developers. We considered those candidates that were called by at least three of the six tools in both the treated and non-treated samples, and not depleted following RNase R enrichment, as true circRNAs (S4 Table). A circRNA candidate is determined to be not depleted if the number of spliced reads per billion mapping (SRPBM; calculated as [number of circRNA supporting reads/total mapped reads] X 10^9^) [2] does not decrease following enrichment. We thus ran ACValidator on these non-depleted potential true circRNA candidates for evaluation.

Overall, except for the SRR1636986-SRR1637090 pair, over 89% of the candidates that were called in both the treated and non-treated pairs were not depleted (i.e., SRPBM after RNase R treatment > SRPBM prior to treatment). Among these non-depleted candidates, ACValidator was able to construct contigs spanning the circRNA junction for more than 75% of them for the RNase R-treated samples and 47-57% of them for the non-treated samples using the medium-stringency criteria and both the window sizes (Table 3). This increased validation rate for the treated samples is expected since RNase R treatment enriches for circRNAs and hence a higher number of back-splice junction supporting reads was observed. Further, higher numbers of validated circRNAs were detected using lower stringency cut-offs for alignment overlap between contigs and junction sequences (Table 3, S4 Table). As observed in the simulation datasets, the different window sizes did not notably affect the number of validations among these experimental datasets (Table 3). Fig 5 shows an example of a circRNA [2,24] that was validated by ACValidator in an MG treated and non-treated pair (reads from this sample aligning to the reference are shown in S1 Fig).

**Table 3.** Summary of ACValidator results on experimental datasets. The “# overlap with pair” column lists the number of circRNAs in common between the RNase R-treated and non-treated samples; “# not depleted” is the number of circRNAs from the overlap whose normalized read counts, i.e., spliced reads per billion mapping (SRPBM) value does not reduce following RNase R-treatment (i.e., SRPBM after treatment > SRPBM before treatment).

| Sample | # overlap<br>with pair | # not<br>depleted | % validated when w = insert size |  |  |  | % validated when w = 2* insert size |  |  |  |
| --- | --- | --- | --- | --- | --- | --- | --- | --- | --- | --- |
|  |  |  | HS | MS | LS | VLS | HS | MS | LS | VLS |
| SRR1636985 | 1,356 | 1,243 | 73.53 | 78.52 | 79.00 | 79.57 | 73.45 | 78.84 | 79.57 | 80.13 |
| SRR1637089 | 1,356 | 1,243 | 46.34 | 49.56 | 50.20 | 51.25 | 44.97 | 49.88 | 50.52 | 51.65 |
| SRR1636986 | 780 | 594 | 79.46 | 84.68 | 85.35 | 86.03 | 80.81 | 86.36 | 87.21 | 87.71 |
| SRR1637090 | 780 | 594 | 48.48 | 54.21 | 55.39 | 55.56 | 48.32 | 55.22 | 56.23 | 56.57 |
| SRR444974 | 953 | 864 | 91.55 | 93.63 | 93.98 | 94.33 | 89.93 | 92.01 | 92.48 | 92.82 |
| SRR444655 | 953 | 864 | 51.74 | 54.98 | 55.21 | 56.60 | 53.36 | 56.94 | 57.18 | 58.91 |
| MG_1 | 1,806 | 1,691 | 72.80 | 77.05 | 77.94 | 78.83 | 71.67 | 77.35 | 78.00 | 78.71 |
| MG_5 | 1,806 | 1,691 | 41.40 | 48.14 | 48.97 | 49.56 | 41.34 | 48.79 | 49.67 | 50.38 |
| MG_2 | 1,430 | 1,331 | 71.90 | 76.86 | 77.46 | 78.14 | 69.72 | 76.11 | 76.63 | 77.31 |
| MG_6 | 1,430 | 1,331 | 40.50 | 47.71 | 48.31 | 49.29 | 40.35 | 47.03 | 47.63 | 48.76 |
| MG_3 | 1,292 | 1,148 | 71.95 | 76.74 | 77.53 | 78.48 | 70.38 | 75.87 | 76.83 | 77.70 |
| MG_7 | 1,292 | 1,148 | 44.08 | 51.48 | 52.26 | 53.40 | 44.51 | 51.39 | 52.18 | 53.57 |

HS, high stringency; MS, medium stringency; LS, low stringency; VLS, very low stringency.

**Fig 5.**
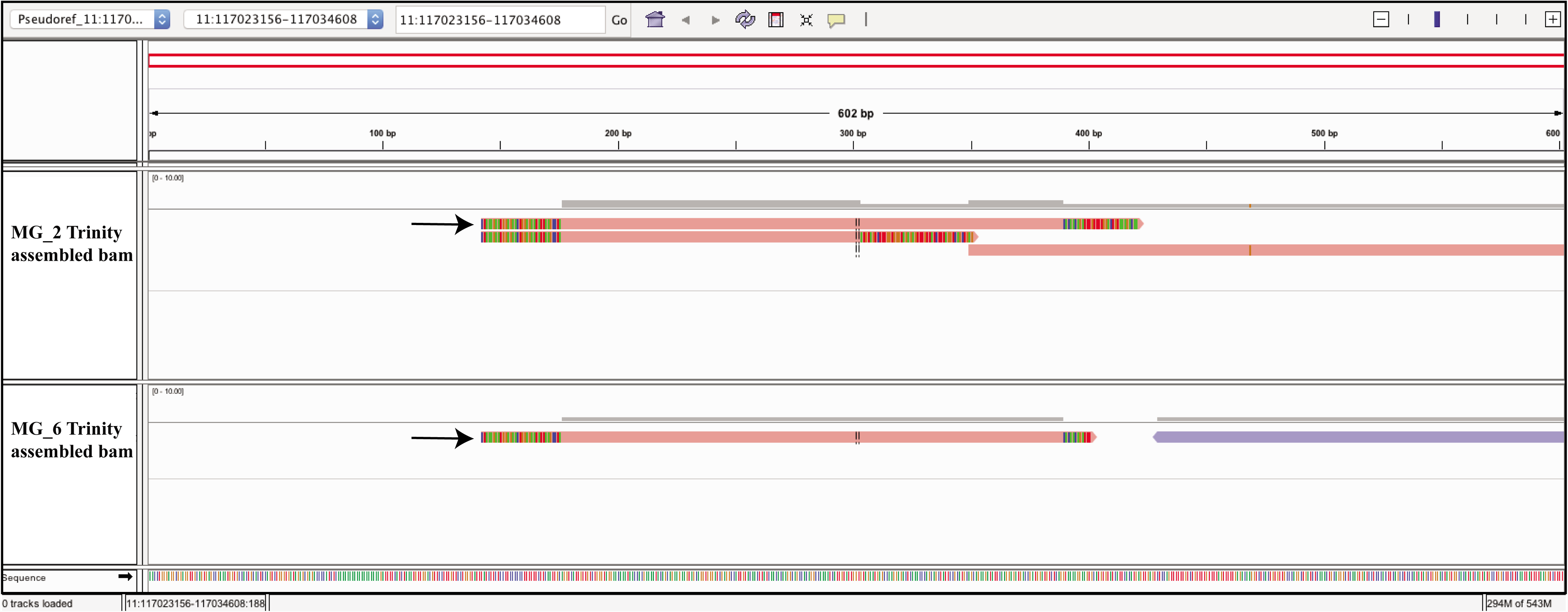
Integrated genomics viewer (IGV) screen shot of a circRNA candidate. (chr11:117,023,156117,034,608) Assembled contigs generated by ACValidator on RNase R-treated (top panel) and non-treated (bottom panel) human middle temporal gyrus (MG) samples, aligned to the corresponding pseudo-reference. This circRNA was detected in both the treated and non-treated samples by at least three of the six existing circRNA prediction algorithms, and was not depleted following RNase R treatment. The circRNA junction of interest is at 300 bp, and pink bars that span over this junction represent the contigs that validate the junction (colored segments at the ends of contigs represent soft-clipped bases; arrows indicate the generated contigs that overlap with the circRNA junction). Reads from these samples aligned to the reference are shown in S1 Fig.

In order to further evaluate the utility of our approach, we next compared the results from each individual tool to those from ACValidator. For this analysis, we used the top 100 most highly expressed candidates from among those we considered as true positives for these experimental datasets (called by at least three of six tools, in both treated and non-treated samples, and SRPBM after RNase R treatment > SRPBM before treatment; S4 Table). Among the in-house treated MG samples, we observed that except for CIRCexplorer and Mapsplice, ACValidator was able to *in-silico* validate a higher number of circRNAs, using the medium stringency criteria (20 bp overlap on either side of circRNA junction), than was detected individually by the other tools (Table 4). Among the SRA samples however, our approach validated a fewer number of circRNAs than were detected by the individual tools except find_circ. Thus, results may vary depending on which individual tool is used for circRNA detection. Notably, the goal of ACValidator is to narrow down a list of potential highconfidence circRNAs and not for comprehensive and *de novo* detection of circRNAs.

**Table 4:** ACValidator and other tools results on top 100 candidates from experimental datasets. The top 100 most highly expressed candidates were selected from the list of circRNAs called by at least three of six tools, in both treated and non-treated samples, and having SRPBM after RNase R treatment > SRPBM before treatment. Each <tool_name>_count column lists the number of circRNAs among the top 100 that were detected by the tool. Similarly, HS, MS, LS and VLS_count columns list the number of circRNAs among the top 100 that were validated by ACValidator using those stringency thresholds.

| Sample | CIRI_<br>count | CIRCexplorer<br>_count | findCirc_<br>count | Mapsplice_<br>count | KNIFE_<br>count | DCC_<br>count | HS_<br>count | MS_<br>count | LS_<br>count | VLS_<br>count |
| --- | --- | --- | --- | --- | --- | --- | --- | --- | --- | --- |
| SRR1636985 | 99 | 93 | 89 | 97 | 92 | 92 | 94 | 97 | 97 | 97 |
| SRR1637089 | 99 | 92 | 87 | 97 | 92 | 92 | 74 | 78 | 79 | 79 |
| SRR1636986 | 99 | 95 | 84 | 96 | 95 | 92 | 91 | 91 | 93 | 94 |
| SRR1637090 | 95 | 88 | 57 | 85 | 95 | 83 | 71 | 77 | 78 | 78 |
| SRR444974 | 99 | 97 | 91 | 97 | 95 | 95 | 97 | 98 | 98 | 98 |
| SRR444655 | 99 | 97 | 76 | 38 | 95 | 90 | 84 | 84 | 84 | 84 |
| MG_1 | 40 | 96 | 85 | 99 | 91 | 88 | 91 | 94 | 94 | 94 |
| MG_5 | 55 | 93 | 72 | 98 | 91 | 80 | 77 | 83 | 83 | 83 |
| MG_2 | 35 | 96 | 90 | 100 | 90 | 90 | 89 | 93 | 93 | 93 |
| MG_6 | 50 | 92 | 68 | 98 | 90 | 82 | 75 | 81 | 81 | 81 |
| MG_3 | 44 | 96 | 83 | 99 | 88 | 88 | 90 | 93 | 94 | 94 |
| MG_7 | 40 | 94 | 79 | 99 | 88 | 85 | 84 | 88 | 91 | 91 |

### PCR validation of identified circRNAs

We performed experimental validations for the top five most highly expressed circRNA candidates that were *in silico* validated by ACValidator and that were detected by all six tools in each sample. Since we were interested in validating the presence of the circRNA and not their abundance, we performed PCR validations on these selected candidates. Each of our treated-non-treated pair is generated from the same donor and hence, we ran validations on the non-RNase R treated cDNA from each donor. For three of the candidates, ACValidator results were experimentally validated (S2 Fig). For the candidate circRNA at chr10:116,879,948116,931,050, ACValidator validated the circRNA junction in two of three samples, while chr9:113,734,352113,735,838 and chr5:38,523,520-38,530,768 were validated in all three samples using all stringency cutoffs. For the remaining two candidates, we observed evidence of validation but because differently sized PCR products were generated, we could not determine the exact product size although it is possible that multiple circRNA species may be present.

### Computational cost overview

We ran our evaluations on a linux-based high-performance computing cluster running CentOS version 7. As expected, the computational cost of our approach directly correlates with the number of reads in the input sample. The only rate-limiting step in using ACValidator is read alignment to generate the SAM file, which is performed prior to starting the workflow. The python validation script following this step requires less than two minutes of runtime for an input SAM file of approximately 8 GB, thus making our approach highly computationally efficient.

### Availability and future directions

We present ACValidator, a novel bioinformatics workflow, which can be used to validate circRNA candidates of interest *in silico* and thus helps to identify true positive candidates. This workflow is applicable to non-polyA-selected RNAseq datasets and can be used to validate circRNAs from various sample types and diseases. When different circRNA detection algorithms identify different circRNA candidates, ACValidator can be used to narrow down specific candidates of interest and thus serve as a circRNA candidate selection tool for experimental validations or functional studies. ACValidator is freely available on our github page: https://github.com/tgen/ACValidator, along with detailed instructions to set up and run the tool (also available as S1 Text).

When evaluated on simulated datasets, ACValidator demonstrates improved performance when higher numbers of circRNA supporting reads are available along with longer read lengths of the sequencing library. When tested on circRNA candidates that were not depleted between RNase R-treated and non-treated sample pairs, we observed a higher validation rate in treated samples, as expected, since those samples are enriched for circRNAs. Window size, the region from where reads are extracted for assembly, is an important parameter for our approach. Through testing of two different window sizes, one equal to and one twice the insert size, we did not observe notable differences in the number of validations. Additionally, we applied different stringency thresholds based on the extent of contig alignment, but we did not observe notable differences in the number of validated candidates across the different thresholds for highly expressed candidates. Although our workflow provides a novel approach for *in silico* validation, we are limited by a few caveats. Primarily, since our assembly analysis relies on reads that extend across circRNA junctions, we are limited in our ability to *in silico* validate circRNAs whose expression may be low, especially in samples that are not enriched for circRNAs. Secondly, since we are limited by the lack of a gold standard circRNA reference dataset, we rely on simulation datasets for evaluation of our approach, which are based on informatically predicted circRNAs detected by various studies and deposited in circBase. Further, since we do not know the true positive events in our experimental datasets, we evaluated candidates that are not depleted by RNase R. However, it is still not known whether RNase R treatment introduces any bias in circRNA detection, especially since some circRNAs are sensitive to RNase R [2,11,25,26].

Future versions of ACValidator will include contig alignment visualization options built into the workflow, as well as alternative strategies to generate contigs for circRNAs with lower expression levels. As circRNAs continue to gain attention as an interesting class of non-coding RNAs, development of novel approaches, including implementation of statistical tests to estimate false discovery rates in circRNA detection, are needed. Continued progress in improving our understanding of the biology of circRNAs will be necessary for such algorithmic development. These findings will be crucial not only for functional analysis, but also for the development of more accurate circRNA detection algorithms.

## Acknowledgements

We are grateful to the Banner Sun Health Research Institute (BSHRI) Brain and Body Donation Program (BBDP) of Sun City, Arizona for the provision of human brain tissues. The BBDP has been supported by the National Institute of Neurological Disorders and Stroke, the National Institute on Aging, the Arizona Department of Health Services, the Arizona Biomedical Research Commission and the Michael J. Fox Foundation for Parkinson’s Research [27]. We also thank Dr. Jonathan Keats, Dr. Sara Nasser, Ryan Richholt (TGen), and Dr. David Craig (USC) for their valuable insights and guidance, Dr. Nancy Linford (Linford Biomedical Communications) for editing assistance, and Andrea Schmitt (Banner Research) and Cynthia Lechuga (TGen) for administrative support.

## Ethics approval and consent to participate

All subjects were enrolled in the BSHRI BBDP in Sun City, Arizona, and written informed consent for all aspects of the program, including tissue sharing, was obtained either from the subjects themselves prior to death or from their legally-appointed representative. The protocol and consent for the BBDP was approved by the Western Institutional Review Board (Puyallap, Washington).

## Supporting Information

**S1 Fig**. Reads from the RNase R-treated (top panel) and non-treated (bottom panel) MG samples, MG_2 and MG_6 respectively, aligned to the human reference genome (hg19).

**S2 Fig. qPCR validation of selected circRNA candidates.** The top five most highly expressed circRNA candidates that were validated by ACValidator and detected by all the six algorithms were selected for validation. Among these, ACValidator was able to validate chr10:116,879,948-116,931,050 in two of the three samples, and chr9:113,734,352-113,735,838 and chr5:38,523,520-38,530,768 in all three samples. Left panel: qPCR of chr5:38,523,520-38,530,768 (junction 1), expected product size: 130 bp; middle panel: qPCR of chr10:116,879,948-116,931,050 (junction 2), expected product size: 679 bp; right panel: qPCR of chr9:113,734,352-113,735,838 (junction 3), expected product size: 76 bp.

**S1 Table: Top 100 most highly expressed circRNAs as well as 100 non-circRNA candidates used for ACValidator evaluation.** The positive and negative candidates used for evaluation from each simulation set are listed in each sheet, which are labeled using the simulation dataset names given in Table 1. Columns D-K indicate whether each circRNA candidate was validated by ACValidator or not, using different stringency criteria and the two window sizes. They are named using the convention “*Validated_<stringency criteria>_<window size>?”.* IS: Insert Size, 2IS: 2 * Insert Size, HS: High stringency, MS: Medium stringency, LS: Low stringency, VLS: very low stringency

**S2 Table: Top 200 most highly expressed circRNAs used for ACValidator evaluation.** The candidates used for evaluation from each simulation set are listed in each sheet along with ACValidator results. Naming conventions are same as in S1 Table.

**S3 Table: Bottom 200 least expressed circRNAs used for ACValidator evaluation.** The candidates used for evaluation from each simulation set are listed in each sheet along with ACValidator results. Naming conventions are same as in S1 Table.

**S4 Table: Non-depleted circRNA candidates in RNase R treated-non-treated pairs (experimental datasets), used for ACValidator evaluation.** The candidates used for evaluation from each sample are listed in each sheet along with ACValidator results. Naming conventions are same as in S1 Table. SRPBM: Spliced reads per billion mapping

**S1 Text: Detailed instructions to set up and run ACValidator.**

