## Supplementary figures and images for "ACValidator: a novel assembly-based approach for *in silico* validation of circular RNAs"

### S1 Fig

#### Supplementary Figure 1

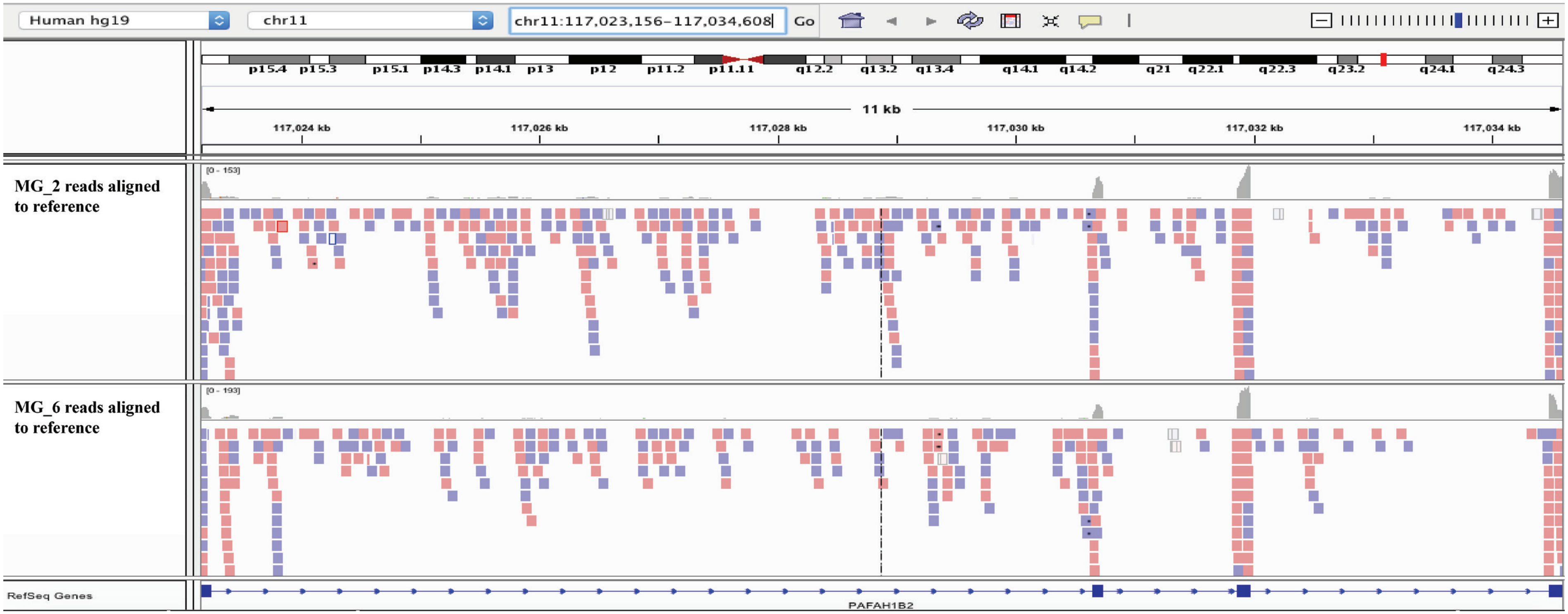

### S2 Fig

Supplementary Figure 2

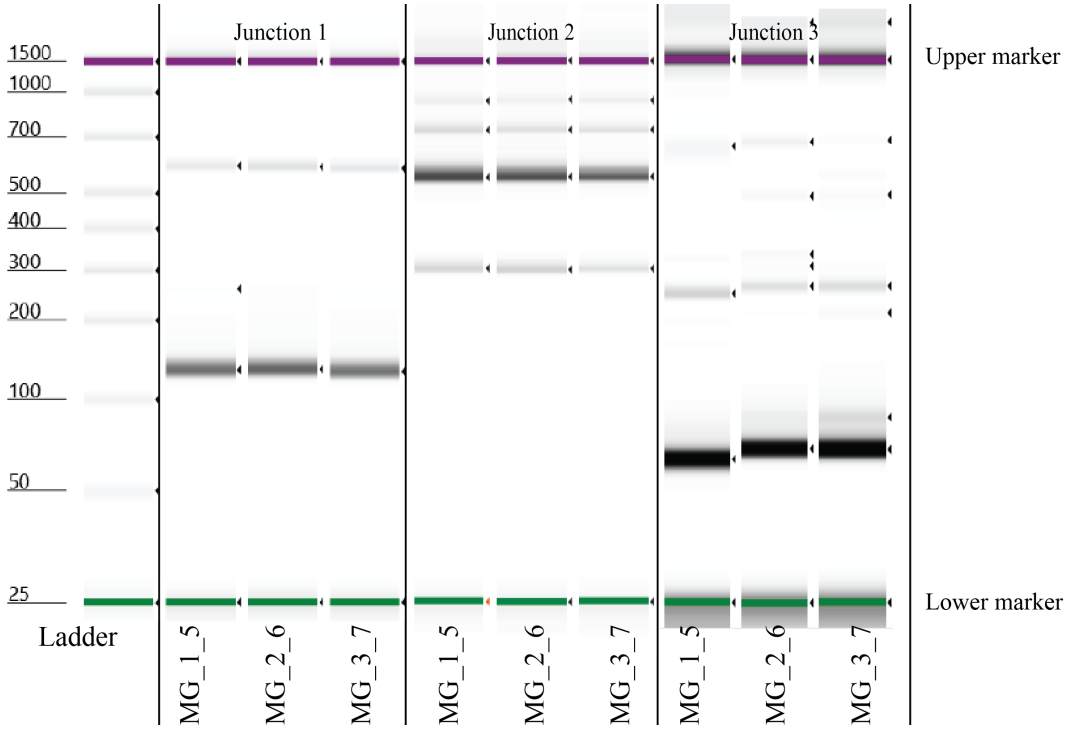
