## Supplementary material for "ACValidator: a novel assembly-based approach for *in silico* validation of circular RNAs": S1 Text

**S1 Text: Detailed instructions to set up and run ACValidator**

All scripts are freely available on our github page: <https://github.com/tgen/ACValidator>

**ACValidator**

Assembly based Circular RNA validator (ACValidator) is a bioinformatics approach to perform in silico validation of selected circular RNA junction(s). ACValidator operates in three phases: extraction of reads from SAM file, generation of a “pseudo-reference” file, and assembly and alignment of extracted reads.

**Set up instructions**

**System Requirements**

Linux OS (tested on CentOS 7)

**Dependencies**

(a) Trinity assembler (tested with v2.3.1) (b) Python with pysam package installed (tested with v2.7.13) (c) Bowtie2 v2.3.0 (d) Samtools v1.4 (e) BWA v0.7.12

There are 2 ways to run ACValidator: 1) directly from source code, or 2) using pip install

**Running ACValidator directly from source code:**

In order to run ACValidator directly using the source code without any installation, copy the main script ACValidator.py and optionally the launcher ACV_launcher.sh (if you have multiple coordinates to validate) to the location of your input SAM file.

Then, to invoke the script for a single SAM file and single coordinate, call as:

*python ACValidator.py -i InputSam -c CircRNA_coordinate -w Window_size --log-filename Log.txt*

To invoke for multiple coordinates, use:

*./ACV_launcher.sh InputSam CoordinateList.txt WindowSize LogFileName.txt*

**Running ACValidator after installation through pip install:**

In order to install ACValidator, clone ACValidator from github and use pip install as below:

*git clone https://github.com/tgen/ACValidator.git cd ACValidator pip install --user .*

Once this is installed, make sure the install location is added in your PATH and then invoke ACValidator from the location of your input SAM file.

To invoke ACValidator for a single coordinate:

*ACValidator -i InputSam -c 8:103312227-103372418 -w 300 --log-filename Log.txt*

To invoke ACValidator for multiple coordinates provided in a text file named CoordinateFile.txt:

*for coordinate in `cat CoordinateFile.txt`; do ACValidator -i InputSam -c ${coordinate} -w ${window_size} --log-filename Log.txt; done*

**Output**

ACValidator results are written to a folder named <InputSam>_validation_tests_<window_size>, which is further organized coordinate wise, with results for each coordinate of interest inside its respective folder. Inside each coordinate folder, ACValidator produces multiple output files, including the sorted sam, bam file of the extracted regions, pseudoreference fasta file, trinity fasta file containing the assembled contigs as well as text files named as “Check_overlap_out_*_coordinate.txt” for each of the 4 stringency criteria. These are 3 column text files containing the name and sequence of the contig that overlaps with the pseudo-reference, as well as the string “Found overlap”. So when these files are empty, or the folder for a coordinate is not created, it means that the respective circRNA coordinate was not validated by ACValidator.

**Notes**

Some circRNA detection tools produce 1-based coordinates while other produce 0-based coordinates. ACValidator assumes that the supplied circRNA coordinates are 0-based. If not, please convert them to 0-based coordinates in order for ACValidator to run properly.

Make sure to change paths in the main script ACValidator.py for dependencies such as Samtools, BWA, Trinity etc to point to the installations in your environment. Also make sure REFPATH folder contains the individual chromosome reference fasta files (e.g., 1.fa, 2.fa...).

**Test data**

<http://tools.tgen.org/Files/ACValidator_test_data/>

Instructions to run are available inside test_data/README_for_testData.txt
